# Inhibiting YAP1 reduced abdominal aortic aneurysm formation by suppressing adventitial fibroblast phenotype transformation and migration

**DOI:** 10.1101/2024.01.02.573973

**Authors:** Cuiping Xie, Yanting Hu, Zhehui Yin, Cuiping Xie

**Author notes:** Correspondence: Dr. Cuiping Xie; Department of General Intensive Care Unit, Key Laboratory of Early Warning and Intervention of Multiple Organ Failure, China National Ministry of Education, Second Affiliated Hospital, Zhejiang University School of Medicine.

## Abstract

**BACKGROUND:** The adventitial fibroblast (AF) is the most abundant cell in the vascular adventitia, a few studies had confirmed that AF contributed to abdominal aortic aneurysm (AAA) development; YAP1 involved in several vascular diseases by promoting AF transformed to myofibroblast, the role of YAP1 in AAA is not clear yet. This study aims to determine whether YAP1 play a role in AAA process by regulating AF function.

**METHODS:** Elastase-induced and Cacl2-induced mice AAA model were used in this study, we applied YAP1 inhibitor-Verteporfin *in vivo* to explore the role of YAP1 in AAA information; *in vitro*, Verteporfin and YAP1-siRNA were used to intervened TGF-β1 induced fibroblasts phenotype transformation and migration model.

**RESULTS:** At first, we use immunohistochemistry (IHC) and western blotting (WB) to compare the expression of YAP1 in aneurysm tissues of AAA patients with normal adjacent vascular tissues, we found that the expression of YAP1 in AAA tissues was significantly increased, and YAP1 mainly increased in adventitia. YAP1 also upregulated in elastase-induced and Cacl2-induced mice AAA model. Then, we diminished YAP1 function with YAP1 inhibitor-Verteporfin in above mice AAA model, AAA incident rate was remarkably declined, and the collagen deposition in the adventitia was alleviated obviously. Afterwards, we studied the effect of YAP1 on the function of AF, we found that the process of AF transforming to myofibroblast and migration were almost completely eliminated after inhibiting YAP1 expression.

**CONCLUSIONS:** This study demonstrated that YAP1 may play a key role in AAA development, inhibiting YAP1 significantly reduced AAA formation through suppressed the process of AF transformed to myofibroblast and migration.

## 1. Introduction

Abdominal aortic aneurysm (AAA) is a chronic inflammatory disease characterized by permanent expansion of local abdominal aortic segments, which is more common in elderly men over 65 years old. Aneurysm rupture is the most serious complication, the mortality rate of which is more than 80%. It is a high-risk vascular disease that leads to an estimated 150,000 to 200,000 deaths worldwide each year^1^.

Previous studies on AAA mainly focused on apoptosis and inflammation of vascular smooth muscle cells, while adventitial fibroblasts (AF), the most abundant cell in the outer membrane of blood vessels, have been ignored. As a receptor of vascular environmental pressure, AF responds to stimulation in a variety of ways, including proliferation, secretion of cytokines, chemokines and growth factors, regulation of extracellular matrix, dilation of nourish blood vessels. By interacting with other cell types, AF coordinates blood vessel responses to injury and stress. AF can also exhibit remarkable plasticity, acquire a smooth muscle cell phenotype (myofibroblast), and participate in vascular remodeling or acquire a pro-inflammatory phenotype through epigenetic changes, thereby recruiting monocytes and macrophages^2, 3^.

Vascular remodeling is the main pathological process of aneurysms, atherosclerosis and stenosis, AF plays a key role in vascular remodeling by transforming into myofibroblasts, which can change the compliance of blood vessels, initiate chronic vascular inflammation, and dilate the nourish blood vessels. Studies have shown that myoblasts of the outer membrane are involved in the pathological processes of abdominal aortic aneurysm, atherosclerosis, vascular stenosis, pulmonary hypertension, and graft vein remodeling^4^. Many factors, such as homocysteinemia, can cause the formation of AAA by promoting AF transformation and the release of pro-inflammatory cytokines^5, 6^。Whether this phenotypic transformation of AF to myofibroblast differentiation plays a protective or promoting role in AAA is still need to be confirmed furtherly, and the specific mechanism involved has not been fully elucidated.

Hippo signaling pathway controls organ size by regulating cell survival, proliferation and apoptosis. YAP1, as a transcriptional coactivator, can bind to TAZ and become a key effector molecule downstream of Hippo signaling pathway. The Hippo/YAP pathway plays an important role in vascular remodeling, influencing the production or degradation of extracellular matrix, migration and death of vascular smooth muscle cells and endothelial cells^7^.

We found that YAP1 expression in the vascular adventitial membrane was significantly increased in both human and mouse AAA tissues. In vitro studies have also confirmed that YAP1 expression is increased during the transformation of AF into myofibroblasts. Previous studies have found that YAP1 can regulate the function of AF through various ways; changes in mechanical stress and hemodynamics can activate YAP1, and subsequently YAP1 regulates the proliferation, phenotypic transformation and migration of AF, thereby causing vascular matrix remodeling^8–10^. The phenotypic transformation of AF is one of the important pathological mechanisms of AAA. Whether YAP1 is involved in this process and ultimately affects the occurrence and development of AAA is still unclear.

## 2. Methods

### 2.1 Human sample and ethics

This study was approved by the Ethics Committee of the Second Affiliated Hospital of Zhejiang University School of Medicine (Approval No. AIRB-2022-06016). All subjects provided informed consent prior to their inclusion in the study, and the experiments complied with the principles outlined in the Declaration of Helsinki.

Human abdominal aortic aneurysm tissues were obtained from AAA patients during the operation, the tissues were fixed with 4% formalin and then embedded with paraffin to prepare tissue microarray sections for immunohistochemical experiments, or stored in liquid nitrogen for subsequent molecular experiments.

### 2.2 Mice

All animal study protocols were approved by the Animal Care and Use Committee of the Second Affiliated Hospital of Zhejiang University School of Medicine (Approval No. IRB-2022-0400), and all experimental procedures complied with the recommendations of the European Ethical Committee (EEC) (2010/63/EU). All animal studies were performed at the Key Laboratory of Early Warning and Intervention of Multiple Organ Failure and Cardiovascular Key Laboratory of Zhejiang Province, Second Affiliated Hospital. 8-10 weeks male C57BL/6 mice were purchased from Shanghai SLAC Laboratory Animal Company. Mice were bred in house under specific pathogen-free conditions with free access to a normal chow diet and water, at a constant temperature (22 ± 2°C) and humidity (60%–65%) with a 12hour dark/light cycle.

### 2.3 Construction of mice abdominal aortic aneurysm model

Elastase-induced and CaCl_2_-induced mice AAA model were used in this study as described previously^11^. Mice were anaesthetized with 2-3%isoflurane, and the infrarenal abdominal aorta was isolated completely, then enfolded with a small piece of gauze soaked in elastase (2.5U/ml, Cat#E1250, Sigma-Aldrich) or CaCl_2_(0.5M, Cat#C5670, Sigma-Aldrich) for 15 minutes, then intra-abdominal was washed with saline, mice were executed under deep isoflurane anesthesia followed by 2 cervical dislocation at the 14 day, the aortas (from the aortic root to the iliac bifurcation) were isolated for embedding or the measurement of aortic diameter or subsequent molecular experiments. To determine the effect of suppressing YAP1on AAA, mice was pretreated with YAP1 inhibitor-Verteporfin (Cat#HY-B0146, MCE) for 24h before the operation in both AAA models, then followed by same research processes mentioned above.

### 2.4 Quantification of abdominal aortic dilation

Incidence of AAA was defined by either (1) 50% or more increase of the maximal diameter in the infrarenal aortic region as compared to the baseline or (2) death due to abdominal aortic rupture^12^. At termination, right atrium was cut open, and PBS was perfused through the left ventricle to remove blood in the aorta. Subsequently, aortas were dissected and placed in 10% neutrally buffered formalin. After fixation, periaortic muscular and adipose tissue was removed thoroughly. Maximal width of infrarenal aortas was measured ex vivo as a parameter for AAA quantification using Image J software. All the data were quantified by two observers that were blinded to the study design.

### 2.5 Isolation and culture of primary AF

Eight-week-old male C57BL/6 mice were selected and executed by deep isoflurane anesthesia followed by 2 cervical dislocation. Sterile instruments were used to expose the abdominal cavity, sterile PBS was fully injected to remove blood in the blood vessels, and the entire abdominal aorta was separated by microforceps. Then the blood vessels were cut open under a microscope, and the media and intima were striked off by microforceps. The adventitial membrane was chopped to the size of 1-2mm^2^ and placed directly into serum-free low-glycemic medium containing 2mg/ml type II collagenase (Cat#17101015, Thermo Fisher Scientific) for digestion for about 4 hours, and beaten every 15 minutes (for 1 minute) until the adventitial membrane was completely decomposed. After digestion, double the volume of serum-containing medium was added to terminate digestion, and after centrifugation at 1500rpm for 15 minutes, the plate was laid. Starting from the third day after the plate was laid, the fluid was changed every 3 days, and the cells were passed when the length reached 80-90%, and the 3-5 generations were used for follow-up experiments.

### 2.6 Wound healing

The AF was plated into the 6-well plate and grew to 100% density, and the 200μl pipette tip was vertically placed at the bottom of the plate to make linear scratch, then washed twice with PBS, and pretreated with Verteporfin(1μM) or YAP1-siRNA(50nM) for 24h, and then recombinant mouse TGF-β1(10ng/ml, Cat#C16W, Novoprotein) was added to continue incubation for 24 hours. Take pictures at 0, 12, 24 hours. The healing area of the scratch is calculated as: migration area (%) =(A0-AN)/A0×100, where A0 represents the initial scratch area and AN represents the scratch area at the shooting time point.

### 2.7 AF phenotypic transformation

AF was plated into 6-well plates, and after being pretreated with (1μM) or YAP1-siRNA (50nM, Zorin bio) for 24h, cells were obtained after incubation with TGF-β1 (10ng/ml) for 24h. The expression of AF to myofibroblast transformation marker-α-SMA was observed by western blotting.

### 2.8 Western blotting

Protein lysate samples were prepared from snap frozen aortas and cells in RIPA solution (Cat# P0013B; Beyotime) supplemented with protease inhibitor (Cat# 05892791001; Roche). Denatured protein lysates were resolved by 8% −10% (wt/vol) SDS-PAGE gels. After transfer, membranes were blocked in 5% (wt/vol) non-fat dry milk diluted in PBS. Membranes were incubated with primary antibodies against YAP1(14074, CST), α-SMA (ab124964, abcam), β-actin (ab197345, abcam) or β-tubulin (ab6046, abcam) overnight at 4°C and subsequently incubated with horseradish peroxidase (HRP) conjugated secondary antibodies which were detected by enhanced chemiluminescence (Cat# WBKLS0500; Millipore). Immunoblots were analyzed using ImageLab software (Bio-Rad).

### 2.9 Statistical analysis

GraphPad Prism 9 was used for statistical analyses. To compare continuous response variables between 2 groups, an unpaired two-tailed Student’s t test was applied for normally distributed variables that passed the equal variance test, and Mann-Whitney U test was used for variables not passing either normality or equal variance test. To compare more than 2 groups, one-way ANOVA and Holm-Sidak method was performed for normally distributed variables that passed equal variance test and Kruskal-Wallis 1 way ANOVA on Ranks with Dunn method for variables failed to pass normality or equal variance test, respectively. P <0.05 was considered as statistically significant.

## 3. Results

### 3.1 YAP1 was increased in the human AAA tissue

We performed immunohistochemical staining (IHC) using tissue microarray sections confirmed that YAP1 was significantly increased in AAA tissues, and was mainly expressed in the adventitial membrane of blood vessels (FIG. 1A). Meanwhile, we compared human AAA tissues to normal blood vessel tissues near aneurysms with western blotting test also showed that YAP1 expression was increased in AAA (FIG. 1B).

**FIG 1:**
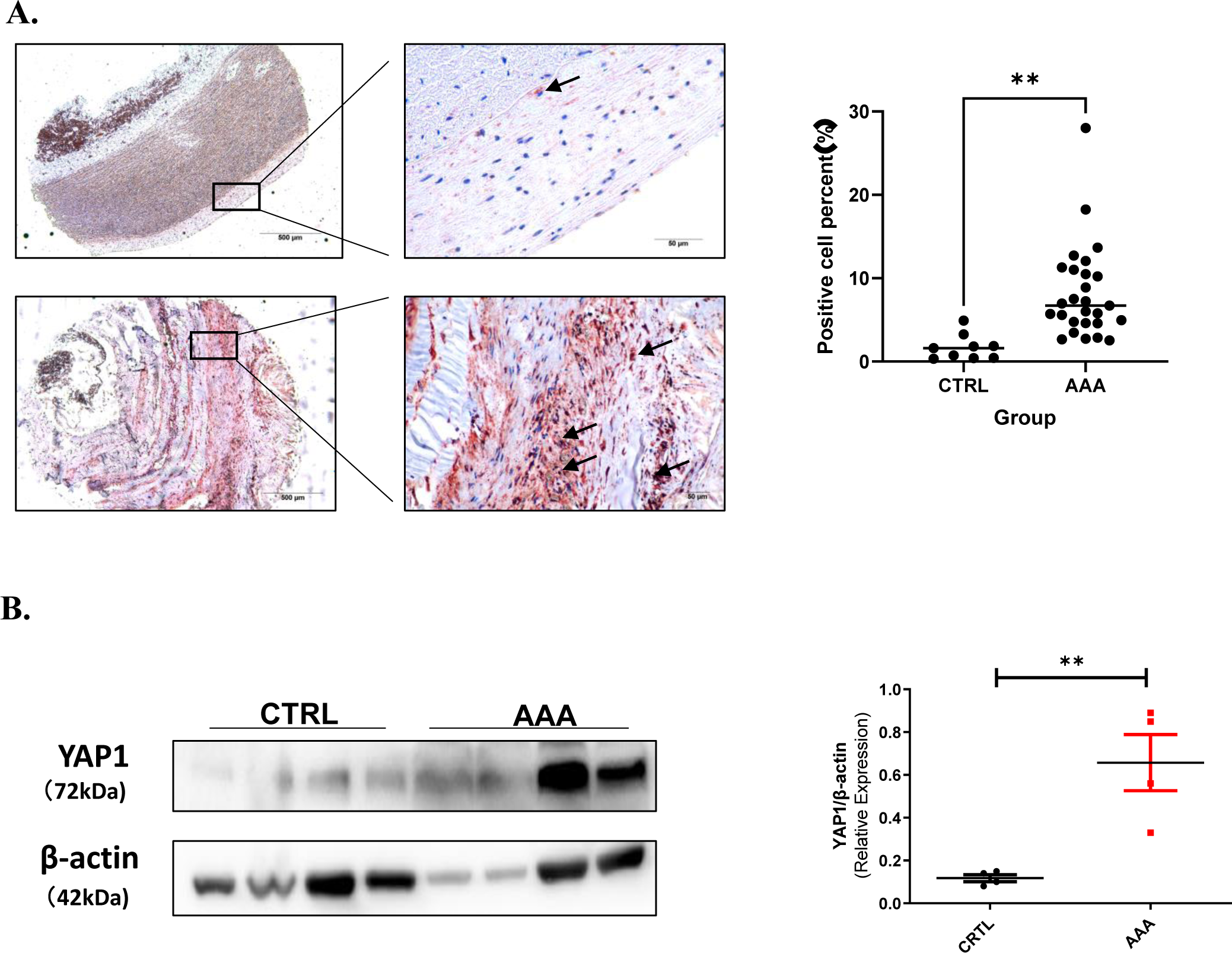
YAP1 was increased in the human AAA adventitial tissue. (A) Representative images of YAP1 expression in aneurysmal lesions(n=26) and normal aorta lesions adjacent to the aneurysm(n=9) as revealed by YAP1 immunostaining using tissue microarray sections; Black boxes indicate the aorta adventitia, black arrows indicate YAP1 positive location; Scale bars represent 500μm(left images) and 50μm(right images), YAP1 positive cell numbers were counted and analyzed with Student *t* test; (B) Protein level of YAP1 in normal aorta lesions adjacent to the aneurysm (n=4) and aneurysmal lesions (n=4), Student *t* test was used for data analysis. Values were represented as mean ± SEM; *P<0.05; **P<0.01; ***P<0.001, respectively.

### 3.2 YAP1 was increased in mice AAA tissues

After the successful establishment of elastase-induced AAA model and CaCl2-induced AAA model, the expression of YAP1 in the control abdominal aorta and AAA tissues was also detected by western blotting and immunohistochemical technique. The expression of YAP1 was increased in the elastase-induced AAA (n=5) tissue compared with that in the control group (saline, n=3) (FIG. 2A); likewise, the expression of YAP1 in the CaCl2-induced AAA (n=5) tissue was significantly increased compare to the control group (saline, n=4) (FIG. 2B). Immunohistochemical staining also confirmed that YAP1 expression was increased in AAA tissues of both models, especially in the adventitial membrane (FIG. 2C).

**FIG 2:**
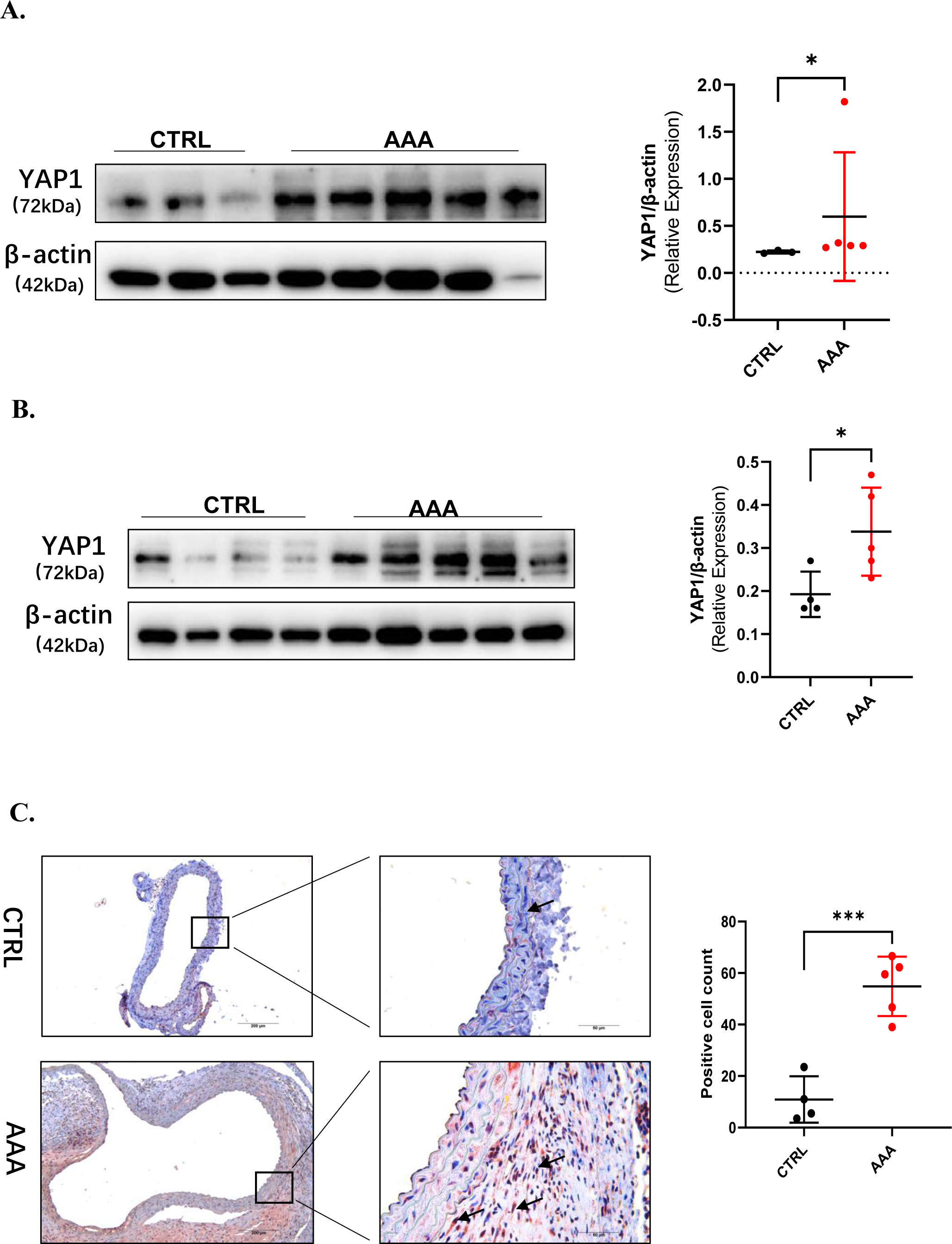
YAP1 was increased in mice AAA tissues. (A) Protein level of YAP1 in control aortas (n=3) and elastase-induced aneurysmal lesions (n=5); (B) Protein level of YAP1 in control aortas (n=4) and Cacl2-induced aneurysmal lesions (n=5); (C) Representative images of YAP1 expression in control aortas (n=4) and elastase-induced aneurysmal lesions (n=5) as revealed by YAP1 immunostaining, scale bars represent 200μm (left images) and 50μm (right images). Black arrows indicate YAP1 positive location. Student *t* test was used for data analysis in (A-C). Values were represented as mean ± SEM; *P<0.05; **P<0.01; ***P<0.001, respectively.

### 3.3 AF phenotypic transformation involved in AAA process

Studies have suggested that phenotypic transformation of AF to myofibroblasts is a significant feature in the pathological process of AAA, which we also confirmed by immunofluorescence staining (FIG 3). α-SMA (-) Vimentin (+) was used as a marker of AF, and α-SMA (+) Vimentin (+) was used as a marker of AF phenotypic transformation (transformation into myofibroblasts).

**FIG 3:**
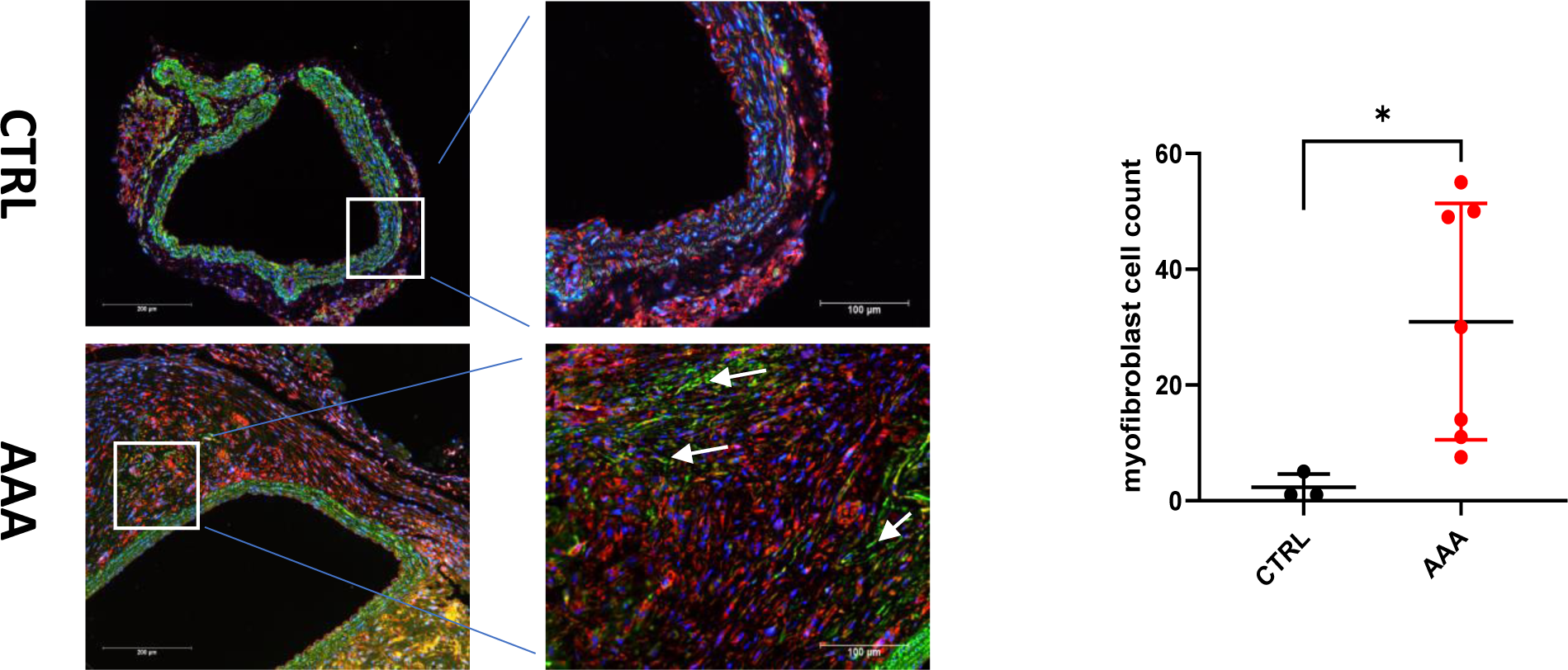
AF phenotypic transformation involved in AAA process. Representative images of α-SMA and Vimentin immunofluorescence in in control aortas (n=3) and elastase-induced aneurysmal lesions (n=7), α-SMA (+) Vimentin (+) was used as a marker of AF transforming into myofibroblasts, myofibroblasts numbers were counted and analyzed with Student *t* test, scale bars represent 200μm (left images) and 50μm (right images), white arrows indicate myofibroblasts location. Values were represented as mean ± SEM; *P<0.05.

### 3.4 YAP1 inhibitor – Verteporfin decreased elastase-induced and CaCl2-induced AAA formation

In order to determine whether YAP1 participates in the formation of AAA, we applied YAP1 inhibitor-Verteporfin to block the function of YAP1 in the elastase-induced and CaCl2-induced AAA model, and observed the abdominal aortic dilation and aneurysm formation rate. Mice were randomly divided into four groups: Control group1 (corn oil + elastase treatment, n=9) and Verteporfin(dissolved in corn oil) group1 (Verteporfin + elastase treatment, n=9); Control group2 (corn oil +CaCl2 treatment, n=9) and Verteporfin group2 (Verteporfin +CaCl2 treatment, n=9); mice in the Verteporfin group were given Verteporfin (30ug/ mouse) by intraperitoneal injection starting from the 24 hours before operation every other day until day 14. We found that in both elastase-induced and CaCl2-induced AAA model, compared with the control group, the abdominal aorta dilatation was significantly reduced in the Verteporfin group (elastase model: 1.515±0.106mm vs 1.072±0.060mm, P<0.01, FIG 4A, B); CaCl2 model 1.615±0.088mm vs 0.815 ± 0.024mm, P<0.001, FIG 4C, D), and the aneurysm formation rate were significantly lower in the Verteporfin groups [elastase model: 22.2%(2/9); CaCl2 model 0%(0/9)] than control group [elastase model: 77.8%(7/9); CaCl2 model 88.9%(8/9)].

**FIG 4:**
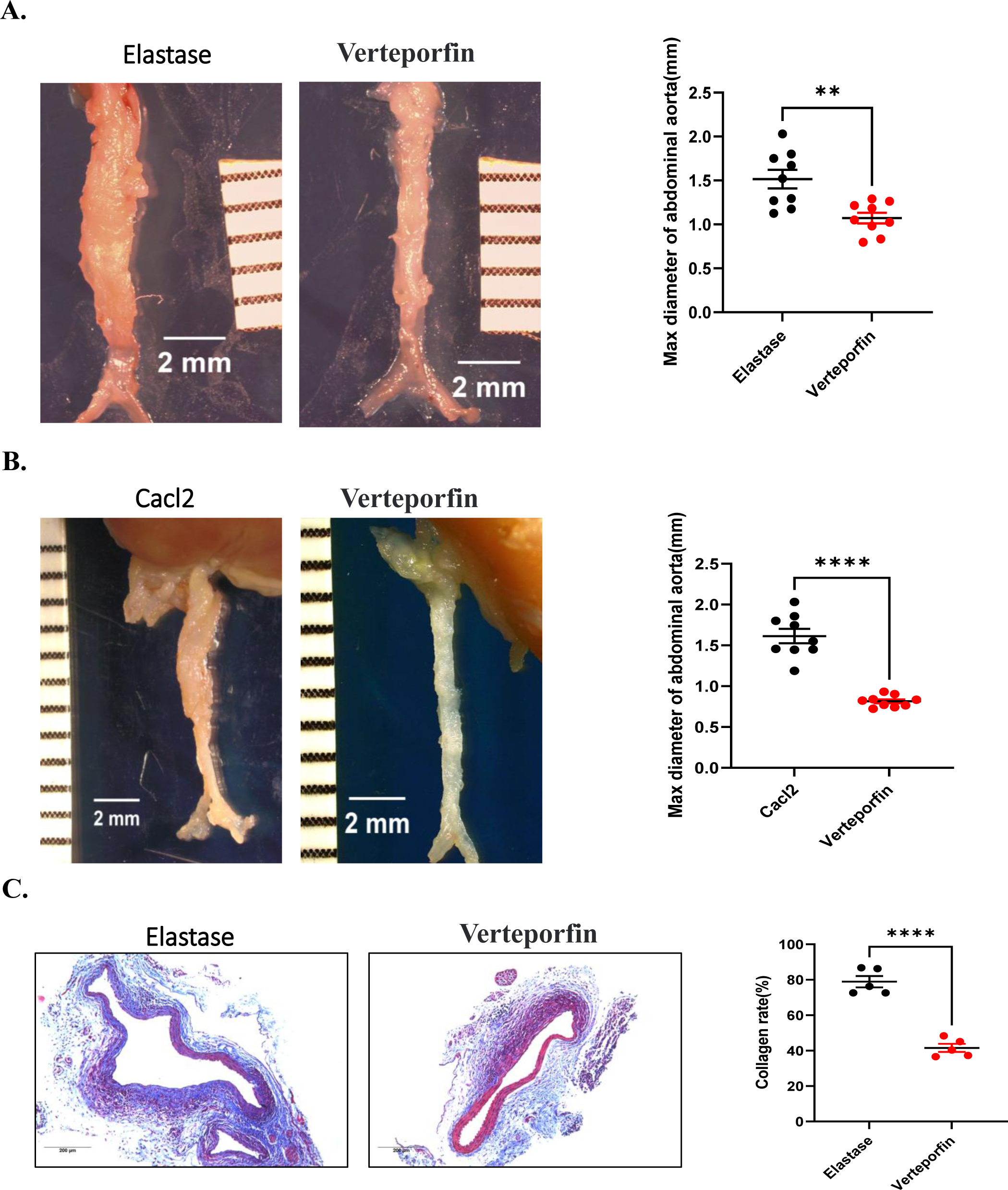
YAP1 inhibitor – Verteporfin decreased elastase-induced and CaCl2-induced AAA formation. (A) Representative images of infrarenal abdominal aortas of elastase-induced AAA (elastase group, n=9) and elastase-induced AAA model pretreated with Verteporfin (Verteporfin group, n=9); (B) Representative images of infrarenal abdominal aortas of Cacl2-induced AAA (Cacl2 group, n=9) and Cacl2-induced AAA model pretreated with Verteporfin (Verteporfin group, n=9). A grid on the scale represents 1mm, scale bars represent 2mm. The ex vivo maximum infrarenal abdominal aorta diameter (external diameter) was measured by image J software, maximum diameter difference between day 14 and baseline in elastase group, Cacl2 group and Verteporfin group were calculated and analyzed with Student *t* test respectively; (C) Representative images of collagen deposition in infrarenal abdominal aortas of elastase group(n=5) and Verteporfin group(n=5) demonstrated by Masson trichrome stain(blue present collagen), scale bars represent 200μm, the collagen rate was calculated by image J software and analyzed with Student *t* test. Values were represented as mean ± SEM; **P<0.01; ****P<0.0001 respectively.

### 3.5 Verteporfin attenuated collagen deposition in AAA adventitia

Collagen deposition is a major pathological feature of AAA, to assess the effect of Verteporfin on collagen deposition, Masson trichrome stain experiment was performed to compare the collagen rate in abdominal aortas from mice AAA model and Verteporfin treated mice. It is showed that the collagen rate is much lower in Verteporfin group (41.6±2.3% vs 78.9±3.2, P<0.001, FIG 4C).

### 3.6 Verteporfin and YAP1-siRNA inhibited AF phenotype transformation and migration

After isolation and culture of AF in vitro, we used TGF-β1(10ng/ml) to construct the cell model of AF transforming to myofibroblast phenotype and migration. Then we pretreatment AF with Verteporfin(1μM) for 24 hours, followed by incubated with TGF-β for 24 hours, α-SMA was used as a marker of AF phenotypic transformation and detected by western blotting; wound healing was performed to assess AF migration; we found that AF transformation and migration were significantly inhibited by Verteporfin (FIG 5A, C). Suppressed YAP1 with YAP1-siRNA also blocked the phenotype transformation of AF (FIG 5B).

**FIG 5:**
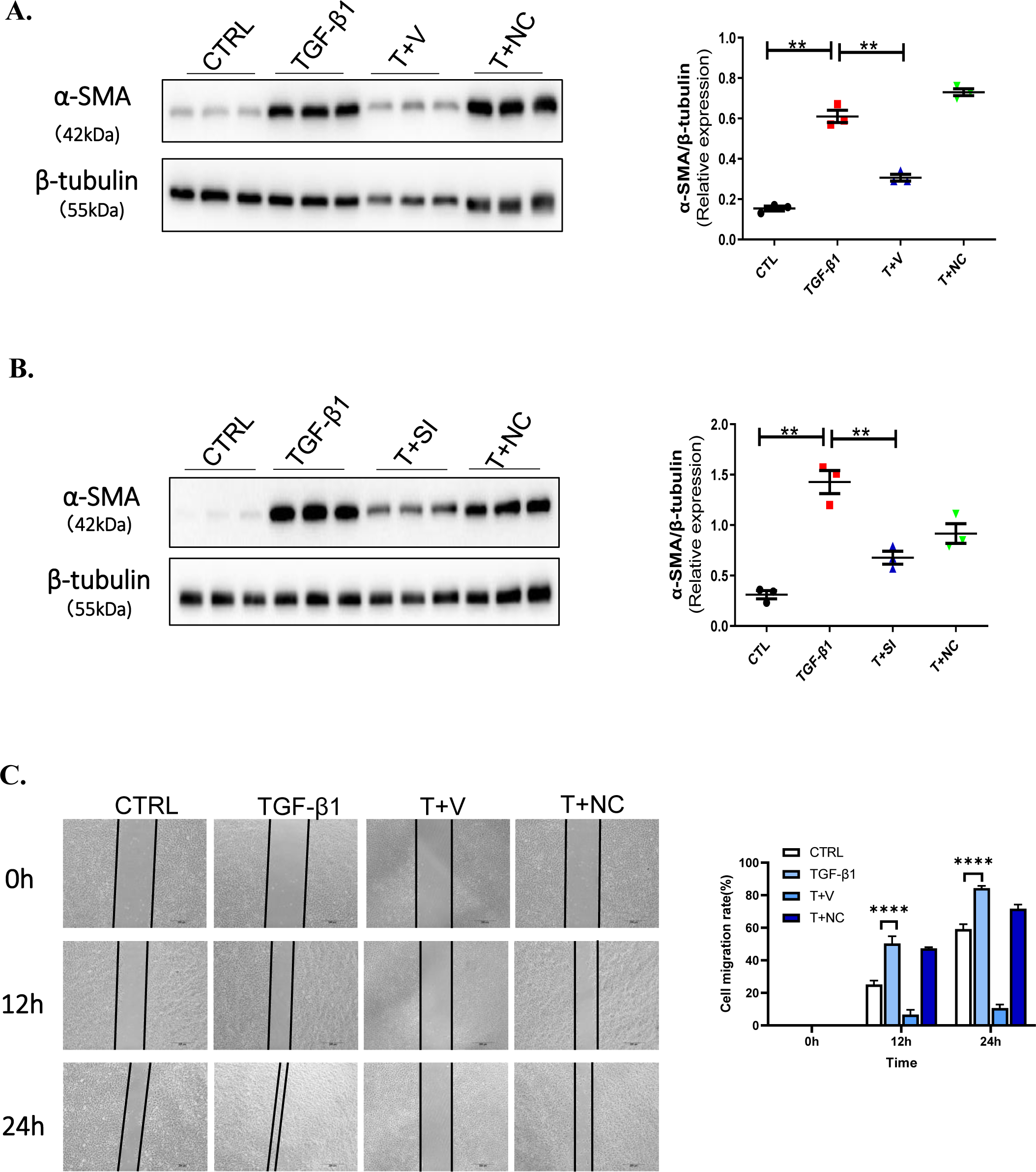
Verteporfin and YAP1-siRNA inhibited AF phenotype transformation and migration. (A) Protein abundance of α-SMA in AF incubated with TGF-β1(10ng/ml) and pretreated 24 hours with Verteporfin (1μM, dissolved with 0.1%DMSO) or negative control (NC, 0.1%DMSO). (B) Protein abundance of α-SMA in AF incubated with TGF-β1(10ng/ml) and pretreated 24 hours with YAP1-siRNA(50nM) or negative control (NC, scramble siRNA, 50nM). (C) Representative images of migration distance of AF incubated with TGF-β1(10ng/ml) and pretreated 24 hours with Verteporfin (1μM, dissolved with 0.1%DMSO) or negative control (NC, 0.1%DMSO) respectively. Scale bars represent 200μm. Values were represented as mean ± SEM; One Way ANOVA was used for data analysis in(A-C), **P<0.01; ****P<0.0001, respectively. T represent TGF-β1, V represent Verteporfin, NC represent negative control (0.1%DMSO in A, C and scramble siRNA in B), SI represent siRNA.

## 4. Discussion

In this study, we used two mice models of AAA-elastase-induced and CaCl2-induced AAA model, demonstrated that YAP1 promoted the development of AAA, and blocking YAP1 in vivo by YAP1 inhibitor can significantly reduce the incidence of AAA and collagen deposition. The main mechanism by which YAP1 affects AAA may be through promoting the transformation of AF into myofibroblasts and migration.

At present, studies on the adventitia of AAA have mainly focused on inflammatory cell infiltration, there are a few studies on AF, the role of AF in AAA has not been clarified. The adventitia of the vessel was considered as a scaffold until recently, when studies confirmed that the adventitial plays a complex and essential role in various vascular diseases. As the most abundant cell in adventitia, AF is the first sensor to environmental stimuli around vascular. Once activated, multiple response such as proliferation, secretion of cytokines and growth factors, alteration of extracellular matrix and expansion of vasa vasorum can occur^13^. In addition, AF has significant plasticity, can be transformed into myofibroblasts, participate in vascular remodeling, secrete inflammatory factors, transform into pro-inflammatory type, and recruit monocytes and macrophages^14^, which also involved in the pathologic process of AAA^5^. Hyperhomocysteinemia (HHcy) promoted Ang II-induced AAA information, it is showed that proinflammatory IL-6 and MCP-1 were colocalized with AF in HHcy and Ang II mice, Hcy sequentially stimulated AF transformation into myofibroblasts, confirmed the essential role of AF in AAA; cylindromatosis (CYLD) also aggravated CaPO4-induced AAA by inducing AF activation(secret proinflammatory cytokines) and transformation^11^. Consistent with previous studies, we found AF transformation contributed to AAA development, and YAP1 activated AF phenotype transformation and migration, which were remarkably blocked by YAP1 inhibition through YAP1 inhibitor-verteporfin or YAP1 siRNA; more critically, intraperitoneal injection of verteporfin significantly attenuated collagen deposition, alleviated AAA development, reduced AAA incidence, verteporfin might be a very promising clinical drug for AAA therapy.

YAP1 and TAZ are mechanosensitive transcriptional coactivators, respond to substrate stiffness, cell density, and shear stress. YAP1 contributed to various of vascular diseases, such as atherosclerosis^15–18^, hypertension^19^, vascular injury, angiogenesis and aneurysm^20–24^. Studies on the mechanism of YAP1 involved in vascular diseases mainly focus on the endothelial cell (EC) and the vascular smooth muscle cell (VSMC), YAP1 contributed to EC dysfunction in the process of atherosclerosis, EC proliferation in hypertension and angiogenesis; YAP1 also induced VSMC proliferation, migration and phenotype transformation in vascular diseases^25^. The role of YAP1 in aneurysm has not been elucidated clearly. It is suggested that YAP1 expression reduced in patients with ascending aortic aneurysms and VSMCs, which was associated with VSMCs apoptosis and extracellular matrix (ECM) disorders of the media^24^.In this study, we confirmed that YAP1 expression upregulated in human AAA tissues through both western blotting and immunohistochemistry, furtherly, we found YAP1 mainly increased in AF in AAA, so we concentrated upon the influence of YAP1 on AF in following research.

The effect of YAP1 on AF has not been fully illustrated. YAP1 has been studied in pulmonary hypertension; it is suggested that YAP/TAZ activation may play a role in initiating pulmonary vascular ECM remodeling by activating pulmonary adventitial fibroblast proliferation in response to increased pulsatility and shear stress^8^. YAP1 promoted AF proliferation and impaired AF apoptosis in pulmonary arteries^26^, controlled ECM remodeling, consequently initiated pulmonary hypertension^27^. Although the current number of studies is small, it is worth believing that AF also play a key role in AAA^5, 11, 28, 29^. Indeed, our study confirmed that AF transformation and migration involved in AAA lesions, which may be facilitated by YAP1; in vivo study showed that YAP1 inhibitor-Verteporfin diminished AF transformation to myofibroblast and migration; in vitro study proved furtherly that YAP1 inhibitor-verteporfin or YAP1-siRNA remarkably attenuated the process of AF transformed to myofibroblast by inhibiting α-SMA expression and migration demonstrated by wound healing experiment.

Above all, this study exhibited for the first time that YAP1 involved in AAA development as far as we known by regulating AF function, it is trustworthy that YAP1 could be promising drug therapy target for AAA patients. Meanwhile, the specific mechanism of YAP1 influenced AF function should be further in-depth studied.

## Acknowledgement

None

